# Biochemical implementation of acceleration sensing and PIDA control

**DOI:** 10.1101/2024.07.02.601775

**Authors:** Emmanouil Alexis, Sebastián Espinel-Ríos, Ioannis G. Kevrekidis, José L. Avalos

## Abstract

Designing dependable, self-regulated biochemical systems has long posed a challenge in the field of Synthetic Biology. Here, we propose a realization of a Proportional-Integral-Derivative-Acceleration (PIDA) control scheme as a Chemical Reaction Network (CRN) governed by mass action kinetics. A constituent element of this architecture is a speed and acceleration biosensing mechanism we introduce and, subsequently, place within a feedback configuration. Our control scheme provides enhanced dynamic performance and robust steady-state tracking. In addition to our theoretical analysis, this is practically highlighted in both the deterministic and stochastic settings by regulating a specific biochemical process *in-silico* and drawing comparisons with a simpler PID controller.

## I. Introduction

Building well-regulated biochemical systems is essential for Synthetic Biology applications to reach their full potential. However, achieving reliable performance at the molecular level requires overcoming significant hurdles associated with a severe lack of robustness and predictability. This necessitates the design of efficient molecular controllers tailored to the unique characteristics of synthetic biological devices [1]–[3].

Proportional-Integral-Derivative (PID) feedback control [4] has arguably been the protagonist in industrial control applications for decades. This has recently motivated several *in-silico* studies to create biochemical analogues for *in-cell* and *in vitro* control [5]–[11]. Overall, PID biocontrollers have been shown to ensure Robust Perfect Adaptation (RPA) [12] - the biological equivalent to robust steady-state tracking - and to provide an acceptable output profile in terms of transient dynamics and, in some cases, stochastic noise. In terms of experimental implementations, fully *in vivo*/*vitro* realizations of PID control have yet to be reported. However, using a hybrid *in vivo*-*in silico* optogenetic platform, the authors in [5] achieved PID control of nascent RNAs in yeast cells by relying on a computer-simulated molecular controller. In addition, simpler control topologies based on integral or proportional-integral action have been successfully implemented in bacterial and mammalian cells, as well as in cell-free environments for a variety of applications [13]–[17].

Despite its prevalence, PID control might not be convenient or even sufficient for achieving tight regulation of high-order processes [4]. To address this challenge, a powerful variation of this concept was proposed by S. Jung and R. C. Dorf, known as Proportional-Integral-Derivative-Acceleration (PIDA) control [18], [19]. In addition to the “three-term” control action of the former, a PIDA controller includes a fourth term, which, in the ideal case, is proportional to the second (double) derivative of the control error (acceleration action). Therefore, it is sometimes referred to as Proportional-Integral-Derivative-Double Derivative (PIDD/PIDD2/PIDD^2^). Although often more structurally complex, PIDA controllers provide significant flexibility for meeting desired performance specifications, such as over-shoot or settling time constraints. Their effectiveness and superior capabilities compared to PID controllers have been extensively investigated in numerous technological applications [20]–[26]. Nevertheless, their utilization in molecular regulation strategies remains unexplored.

In this work, we first introduce a simple method of acceleration sensing at a molecular level by leveraging the mechanism of the *Biomolecular Signal Differentiator-II* (*BioSD-II*) module [27]. Specifically, we present a single-input two-output biochemical device, termed *BioSD*^2^, which is capable of estimating the first and the second derivative of a molecular signal. Additionally, *BioSD*^2^ exhibits strong (low-pass) filtering action, offering protection against undesired amplification of high-frequency input signal components - a major obstacle in signal differentiation applications [4], [27]. We also introduce the biorealization of a PIDA controller equipped with some advanced features (compared to its “textbook” version) and discuss a few control variations arising under different structural conditions. Subsequently, we practically demonstrate the advantages of our control scheme by considering a biochemical (open-loop) process of three, mutually interacting species and providing a comparative computational analysis in terms of the (closed-loop) deterministic and stochastic performance based on PIDA and PID control. Note that all the biochemical topologies in this work are represented via the widely adopted formalism of Chemical Reaction Networks (CRNs), following the law of mass action [10], [28].

The remainder of the paper is organized as follows: Section II discusses important background principles; Sections III and IV introduce the CRN implementation of *BioSD*^2^ and the PIDA controller, respectively; Section V presents numerical simulation results; and Section VI concludes our work and outlines future research directions.

## II. Background

### A. PIDA control

Fig. 1(a) illustrates an ideal PIDA control algorithm described by:

**Fig. 1:**
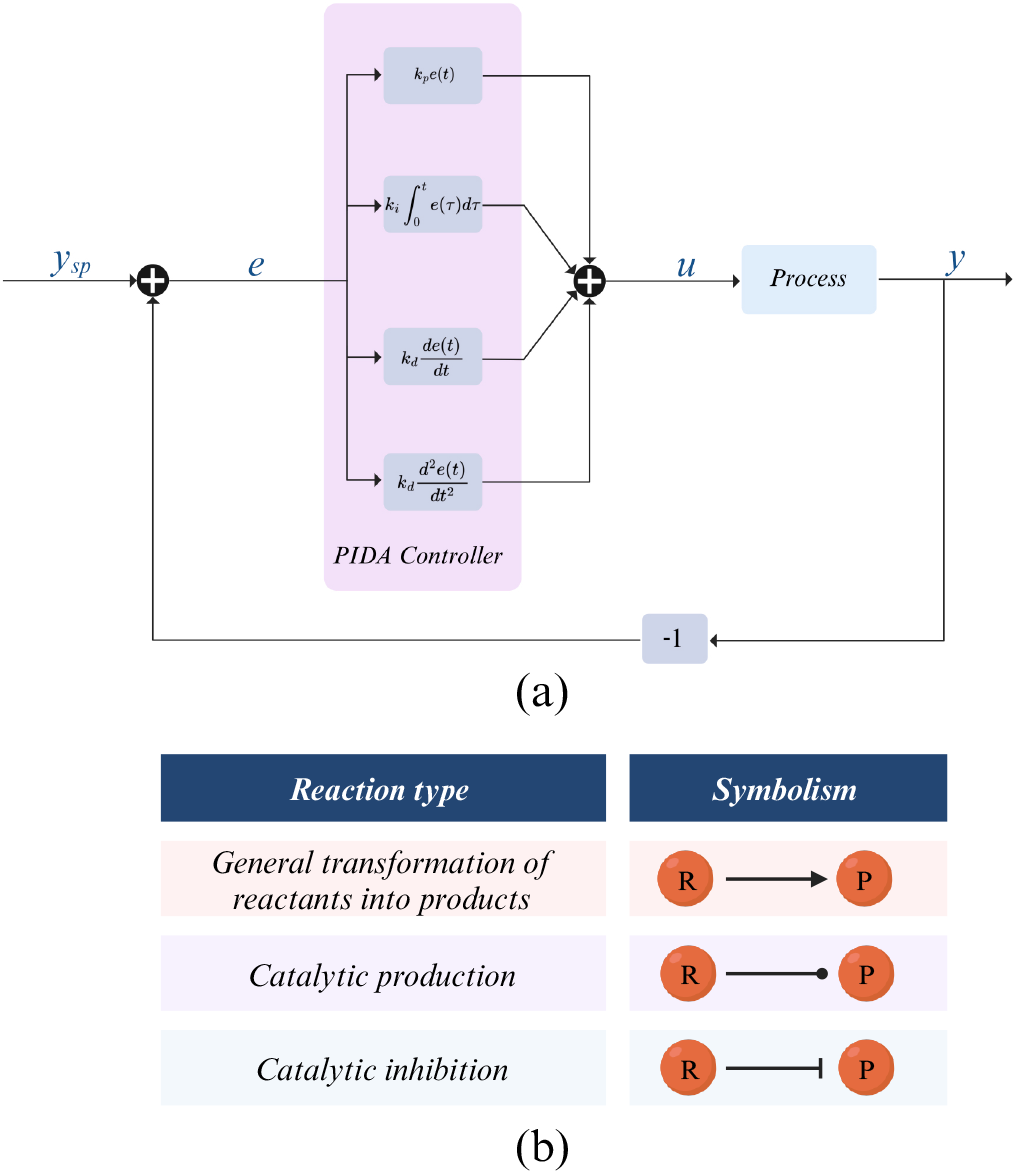
(a) An ideal PIDA control scheme based on error feedback. (b) Biochemical interactions employed in this study [29].

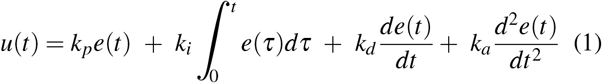

with *e*(*t*) = *y*_*sp*_ − *y*(*t*) and *u*(*t*), *y*(*t*), *y*_*sp*_, *e*(*t*) representing the control input, process output, set-point and control error, respectively.

Equation (1) is often encountered in the Laplace domain, where it can be written as:

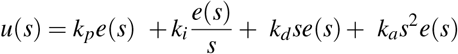

where *s* is the Laplace variable (complex frequency) and *u*(*s*), *e*(*s*) represent the Laplace Transform of *u*(*t*), *e*(*t*), respectively.

This type of control schemes can be further refined to achieve enhanced performance [4], [20]–[22], [30]. For example, to deal with high-frequency noise amplification resulting from signal differentiation, a low-pass filter can be embedded into the last two terms of Equation (1). Simultaneously, replacing 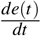 and 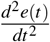 in the latter terms with 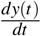 and 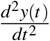, respectively, can eliminate the so-called *derivative* and *acceleration kicks*, i.e. undesired transients of the control signal due to sudden set-point changes. Additionally, *k*_*p*_*y*(*t*) is often favored over *k*_*p*_*e*(*t*) (in Equation (1)) to further improve transient dynamics by eliminating the *proportional kick*. It is worth noting that replacing the error with the output signal regarding proportional, derivative and acceleration actions constitutes a special case of *set-point weighting*. The above adjustments are also adopted in this work.

### B. Modelling and numerical simulations

The biochemical architectures herein are composed of the following types of mass-action-kinetics-based chemical reactions (Fig. 1(b)) [28], [29]: *R* ⟶ *P* (general transformation of reactants into products), *R* ⟶ *R* + *P* (catalytic production), *R* + *P* ⟶ *R* (catalytic inhibition), where *R, P* are two biochemical species.

The analysis in this work employs deterministic mathematical models based on Ordinary Differential Equations (ODEs). This approach typically offers a good approximation of a system’s dynamics, provided that the concentrations of the involved species are sufficiently high. In parallel, to investigate stochasticity, we utilize Gillespie’s stochastic simulation algorithm (SSA) [31]. The statistics presented are derived from 5 *·* 10^4^ stochastic trajectories (unless otherwise stated). All numerical simulations were performed using BasiCO [32], a Python interface to the biochemical system simulator COPASI [33].

## III. Speed and Acceleration biosensing

Here we introduce a biochemical device referred to as *BioSD*^2^, with one input, *U*, and two outputs, *X*_1*S*_ and *X*_1*A*_, whose CRN architecture is illustrated in Fig. 2(a). *BioSD*^2^ can be seen as the “catalytic production interconnection” of two *BioSD-II* modules with no “self-degradation” of the output species, as expounded in [27]. The first *BioSD-II* module encompasses species *X*_1*S*_, *X*_2*S*_, *X*_3*S*_ while the second one comprises *X*_1*A*_, *X*_2*A*_, *X*_3*A*_.

**Fig. 2:**
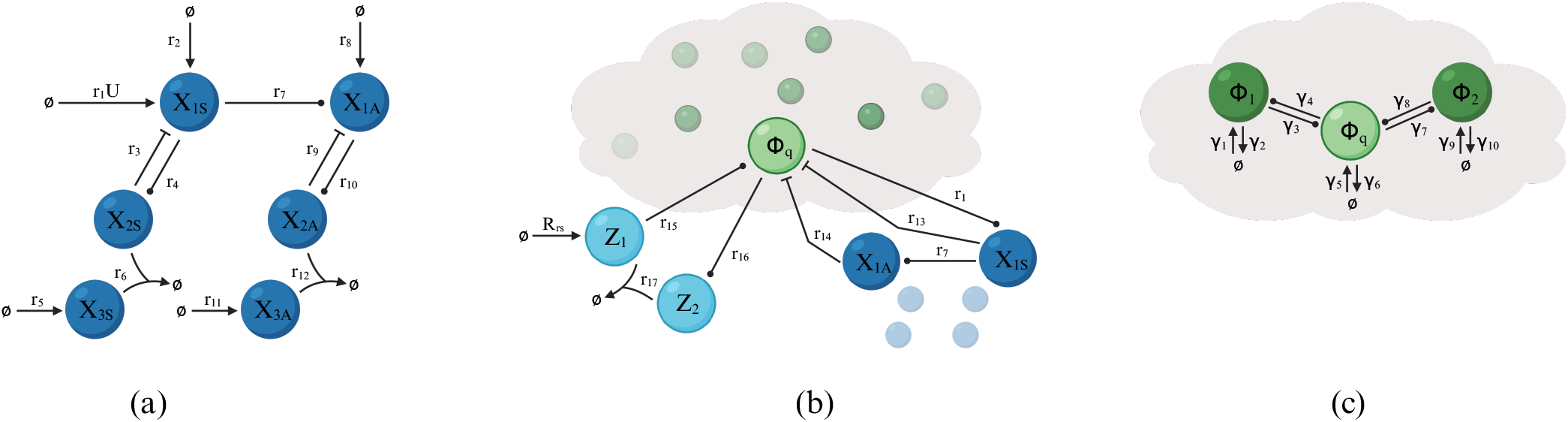
Chemical Reaction Network (CRN) implementation of (a) the *BioSD*^2^ module (Section III) (b) a PIDA feedback control scheme (blue color) for a general biochemical process to be controlled (green color) - cloud network (Section IV). For visualization purposes, only the part of *BioSD*^2^ that directly interacts with the cloud network is shown in detail, with the remainder represented by small, faint circles. (c) a three-species biochemical process as the network to be controlled (cloud network) (Section V).

The dynamics of *BioSD*^2^ is given by the ODE model:

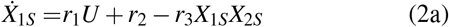

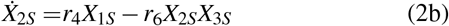

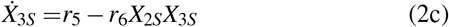

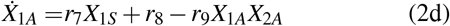

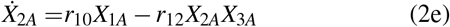

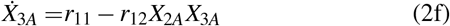

where *r*_*i*_ ∈ ℝ_+_ with *i* ∈ ℕ and 1 ≤ *i* ≤ 12.

Taking into account the analysis in [27], it is straightforward to show that, for any constant input, *U*^*^, system (2) has a unique positive steady state 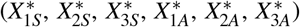, which is locally exponentially stable. In addition, assuming sufficiently small perturbations *u, x*_1*S*_, *x*_2*S*_, *x*_3*S*_, *x*_1*A*_, *x*_2*A*_, *x*_3*A*_ around this fixed point and:

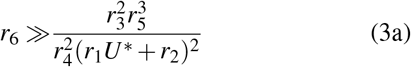

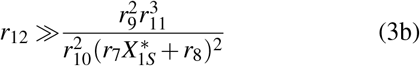

the corresponding input/output relations near the steady state are described in the Laplace domain as:

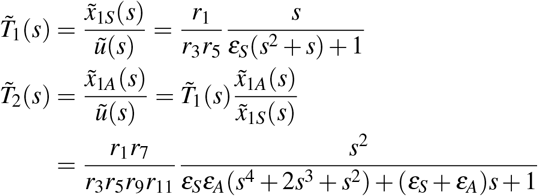

where 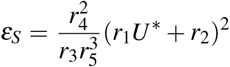 and 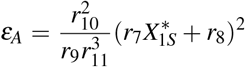, and 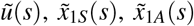 represent the Laplace Transform of *u, x*_1*S*_, *x*_1*A*_, respectively. Moreover, constraints (3a) and (3b) refer to the *BioSD* modules described by Equations (2a)-(2c) and (2d)-(2f), respectively. As demonstrated in [27], these constraints arise from a singular perturbation analysis of the corresponding equations, ensuring that the first derivative of a molecular input signal is (locally) estimated with high accuracy.

As 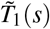 and 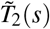 indicate, the speed (first derivative) and the acceleration (second derivative) of *u* are estimated via *x*_1*S*_ and *x*_1*A*_ accompanied by the action of a second- and fourth-order low pass filter, respectively. These filters can be tuned to satisfy the performance specifications of interest by adjusting *ε*_*S*_ and *ε*_*A*_ - more details on this topic can be found in [27].

Transitioning to the time domain [27], if an input signal is (sufficiently) slow and of a (sufficiently) long time duration, the outputs of *BioSD*^2^ can be locally approximated as follows:

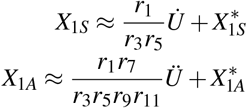

where 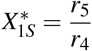 and 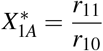.

## IV. Biorealization of PIDA control

We now present our PIDA bio-controller, with its CRN architecture depicted in Fig. 2(b). We consider a general biochemical process to be controlled (cloud/open-loop network) with Φ_*q*_ representing the target species (output species of interest) and being free to participate in an arbitrary number of reactions. Although detailed knowledge of the cloud network’s structure (species/reactions) is not necessary, accessibility of Φ_*q*_ to the controller is a requirement. To achieve a minimal structural design, we establish the actuation and sensing processes between the two. Within the controller’s architecture, two primary building elements can be observed: an *antithetic integral motif* [34], collectively formed by species *Z*_1_, *Z*_2_, and a *BioSD*^2^ device, as discussed in Section III. It is important to emphasize that our analysis is restricted to cases where the existence of an asymptotically stable and biologically meaningful equilibrium of the closed-loop system can be guaranteed.

Given a cloud network with *q* species, Φ_1_, Φ_2_,…, Φ_*q−*1_, Φ_*q*_, the closed-loop dynamics can be captured by the ODE model:

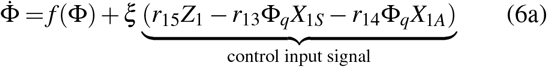

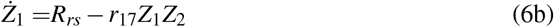

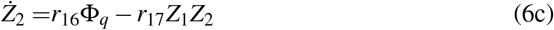

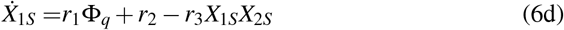

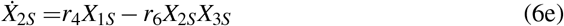

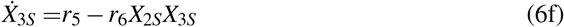

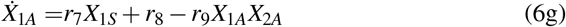

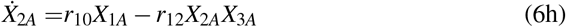

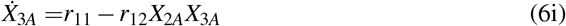

where *R*_*rs*_ corresponds to a non-negative, time-varying reference signal and *r*_*i*_ ∈ ℝ_+_ with *i* ∈ ℕ and 1 ≤ *i* ≤ 17. The term *f* (Φ) represents the dynamics of the open-loop process, Φ = [Φ_1_ Φ_2_ … Φ_*q−*1_ Φ_*q*_]^*T*^ and *ξ* = [0 0 … 1]^*T*^ ∈ ℤ^*q*^.

Assuming an equilibrium of interest for the closed-loop system, denoted as *E*, achieved for some constant reference signal, 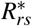, we shift our focus on the (closed-loop) behaviour near this equilibrium. We therefore consider coordinate transformations of the form *ψ* = Ψ *−* Ψ^*^ which represent sufficiently small deviations around this point. Here, Ψ and Ψ^*^ respectively refer to any variable of system (6) and its associated steady state. Subsequently, applying Jacobian linearization yields:

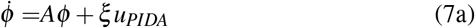

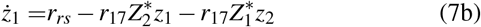

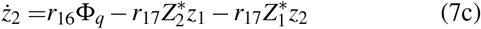

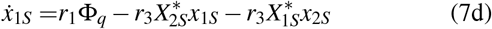

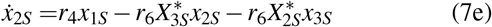

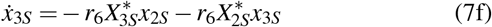

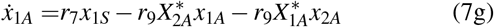

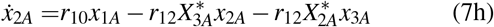

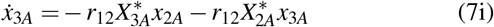

where 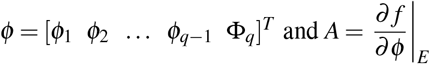. Moreover, for the control input signal we have:

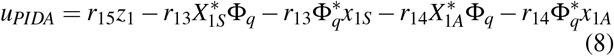

Equations (7c), (7b) yield:

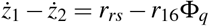

Or

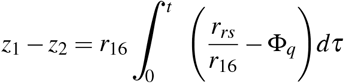

which reveals the robust steady-state tracking property due to the presence of integral action. The error signal can therefore be defined as 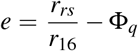, where 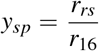 and *y* = Φ_*q*_ according to the notation in Section II.A.

Moving to the Laplace domain, Equation (8) can be expressed as:

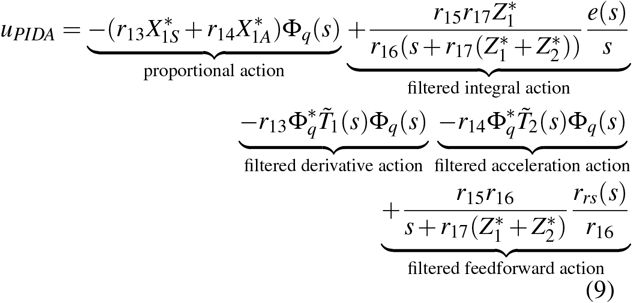

where 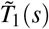 and 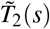 are defined in Section III. Note that, to align with the analysis presented in this section, the term *U*^*^ appearing there must be replaced by 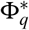.

Assuming now that *antithetic integral motif* operates in the fast sequestration regime, i.e., *r*_17_ → ∞, [5], we get:

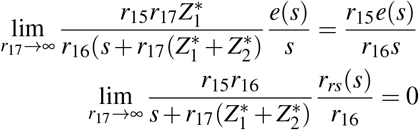

since 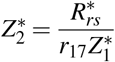 and 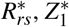 are independent of *r*_17_. This can be easily seen from system (6) if we consider that *f* (Φ) refers to the cloud network and 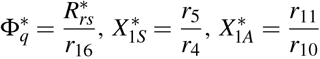.

Thus, the control law (9) can be written as:

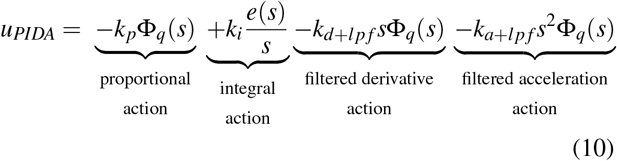

Where 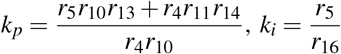,

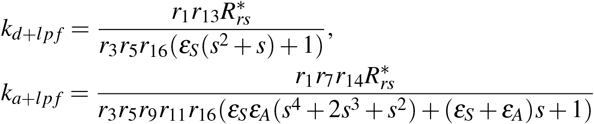

with 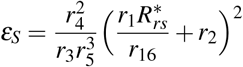 and 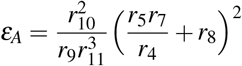.

Furthermore, the inequalities (3) become:

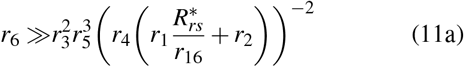

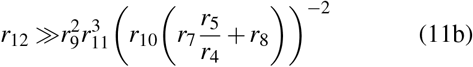

As shown above, the integral term in Equation (10) is realized via the *antithetic integral motif*, while the remaining terms are realized via the *BioSD*^2^ module. In applications that do not require precise output regulation, the former topology can be omitted, resulting in a simplified PDA control law, which can be seen as an extension of PD control schemes. Note that a multicellular PD controller was recently proposed in [35].

Achieving closed-loop stability is an essential requirement for the proposed control scheme, which can be ensured by selecting appropriate parameter regimes for the reaction rates involved (assuming that such regimes can be practically achieved). The tuning process can be facilitated by computing the Jacobian matrix of system (6). If this matrix is Hurwitz, then the local asymptotic stability of the equilibrium of interest can be guaranteed. To determine necessary and sufficient conditions for this, one can use the Routh-Hurwitz criterion.

## V. Regulating a three-species biochemical process

In Fig. 2(c) we present a biochemical process involving three species with mutual interactions. This replaces the abstract cloud network in the closed-loop system shown in Fig. 2(b). The resulting closed-loop behaviour is given by:

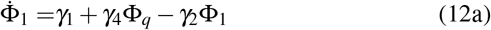

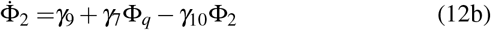

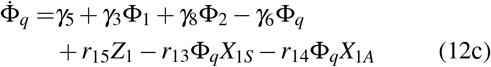

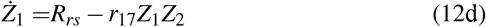

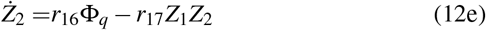

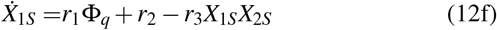

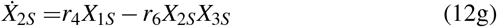

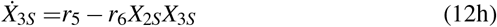

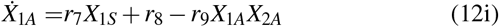

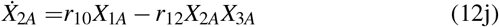

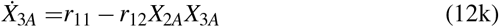

with *γ*_*m*_, *r*_*i*_ ∈ ℝ_+_ with *m, i* ∈ ℕ and 1 ≤ *m* ≤ 10, 1 ≤ *i* ≤ 17.

In what follows, we computationally compare the output response (species Φ_*q*_) in two scenarios: PIDA-based regulation (see Fig. 2(b)) and PID-based regulation (see Fig. 2(b) without taking into account species *X*_1*A*_, *X*_2*A*_, and *X*_3*A*_).

### A. Deterministic simulations

Fig. 3(a) shows the output response in the deterministic setting. The PIDA controller results in a smooth transient response, eliminating the overshoot that appears when the PID controller is used. As expected, in both cases, the output converges to the same steady state due to the presence of the integral control acting on the same error quantity.

**Fig. 3:**
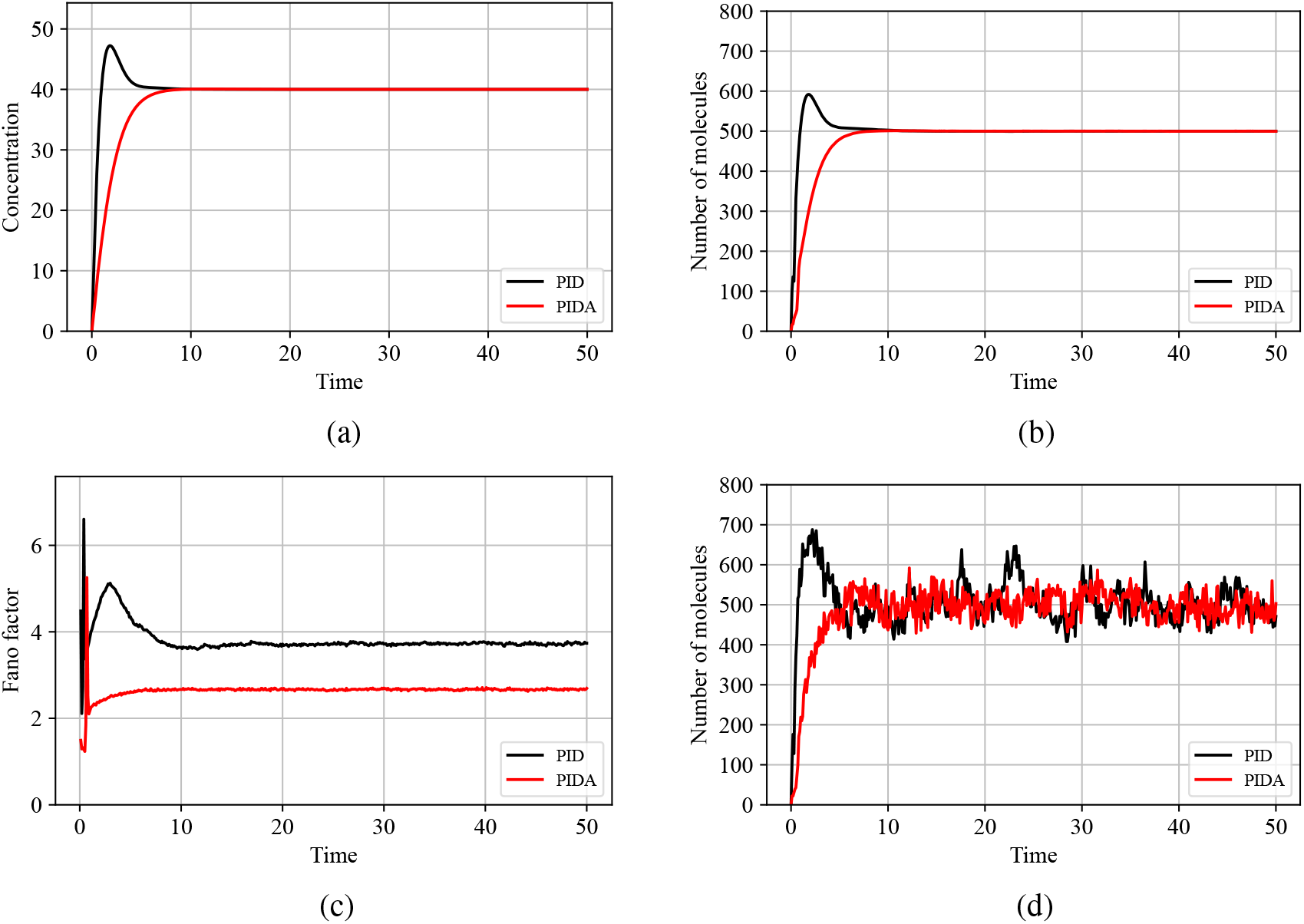
(a) Deterministic setting (Section V.A): Simulated output response (species Φ_*q*_) of system (12) using the following (arbitrarily chosen) parameters: *γ*_1_ = 50, *γ*_2_ = 0.5, *γ*_3_ = 2, *γ*_4_ = 2, *γ*_5_ = 1, *γ*_6_ = 0.5, *γ*_7_ = 3, *γ*_8_ = 1, *γ*_9_ = 1, *γ*_10_ = 0.5, *r*_1_ = 100, *r*_2_ = 150, *r*_3_ = 1, *r*_4_ = 1, *r*_5_ = 100, *r*_6_ = 1000, *r*_7_ = 100, *r*_8_ = 150, *r*_9_ = 1, *r*_10_ = 1, *r*_11_ = 100, *r*_12_ = 1000, *r*_13_ = 0.2, *r*_14_ = 0.9, *r*_15_ = 100, *r*_16_ = 0.5, *r*_17_ = 1000, *R*_*rs*_ = 20. The PIDA case is described by system (12). PID case is described by the same system if we neglect Equations (12i)-(12k) and, consequently, the term *−r*_14_Φ_*q*_*X*_1*A*_ in Equation (12c). Note that the above parameters ensure that the *antithetic integral motif* practically operates in the fast sequestration regime and the constraints (11) are satisfied. Moreover, in both cases, linear asymptotic stability is guaranteed as the eigenvalues of the corresponding Jacobian matrix have negative real parts. Specifically, for the PIDA case, we have: *−*36570.55, *−*101500.99, *−*41502.41, *−*157.66, *−*46.92 *±* 59.62 *j, −*0.8 *±* 0.77 *j, −*0.5, *−*0.45 *±* 0.14 *j*, while for the PID case, we have: *−*41502.41, *−*605.17, *−*29.89 *±* 27.63 *j, −*1.02 *±* 1.14 *j, −*0.61, *−*0.61, *−*0.5. (b)-(d) Stochastic setting (Section V.B): The corresponding (b) stochastic means (c) Fano factors, *F*_*a*_(*t*), and (d) two (individual) stochastic trajectories (one for each case) are computed using SSA. The conversion of concentration (deterministic setting) to the number of molecules (stochastic setting) was made using the relation *n*_*m*_ = *cVN*_*A*_, where *n*_*m*_ is the number of molecules, *c* is the concentration, *V* is the reaction volume and *N*_*A*_ is Avogadro’s constant. The value for *V* was selected arbitrarily.

### B. Stochastic simulations

The above behaviour is also observed in terms of the stochastic mean of the output responses (see Fig. 3(b)).

Additionally, as a metric to quantify the output noise, we use the Fano factor [36] defined as:

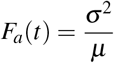

with *σ* and *µ* representing the standard deviation and mean, respectively. Overall, PIDA control yields a considerably smaller *F*_*a*_(*t*) (see Fig. 3(c)) and, consequently, less noisy individual stochastic trajectories (see Fig. 3(d)) compared to PID control. This aligns with the noise-reduction capability of BioSD-based feedback action, as demonstrated in other molecular control architectures [7], [11]. It is worth mentioning that noise reduction through (first-order) derivative control action, achieved via different mechanisms, has also been observed [5], [9].

### C. Perturbation of the kinetic parameters

We subsequently perturb all the reaction rates involved in the closed-loop architecture by 25% adopting different simulation scenarios. As shown in Figs. 4, 5, and 6, PIDA control outperforms PID control by generally offering a significantly more robust output performance associated with reduced stochastic noise.

**Fig. 4:**
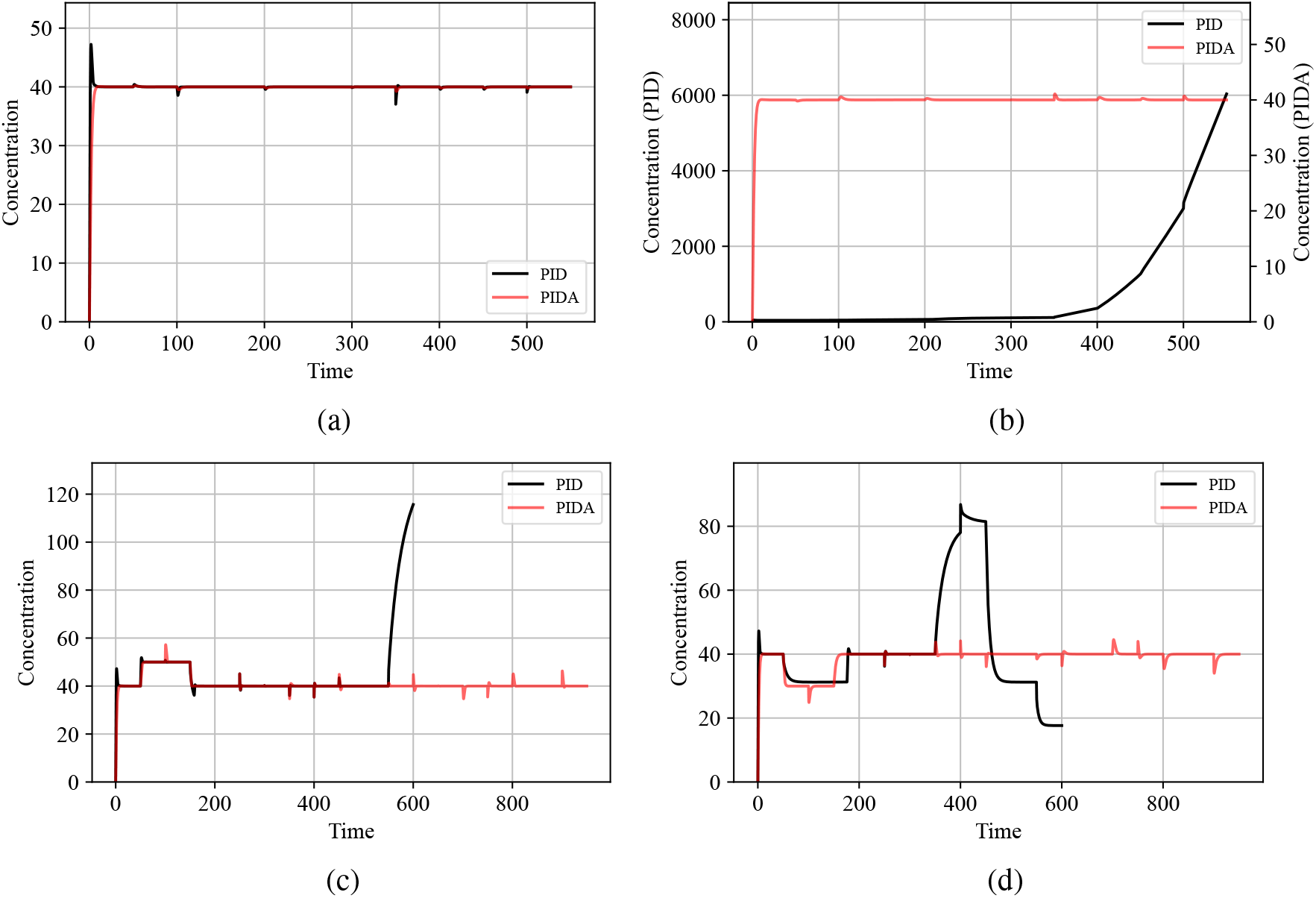
Simulated output response based on the deterministic simulation setup of Fig. 3(a) in the presence of disturbances/ parametric uncertainty. In particular, every 50 time units a perturbation (increase or decrease) of 25% is applied to a different reaction rate in the closed-loop system, according to the following (arbitrarily chosen) scenarios: (a) Increase in *γ*_1_, *γ*_2_, *γ*_9_, *γ*_10_, *γ*_5_, *γ*_6_ and decrease in *γ*_3_, *γ*_4_, *γ*_7_, *γ*_8_, in the order they appear (b) The parameters increased or decreased in scenario (a) are now decreased or increased, respectively, in the same order. (c) Increase in *R*_*rs*_, *r*_15_, *r*_16_, *r*_17_ (associated with the *antithetic integral motif*) and decrease in *r*_1_, *r*_2_, *r*_4_, *r*_3_, *r*_5_, *r*_6_, *r*_13_, *r*_7_, *r*_8_, *r*_10_, *r*_9_, *r*_11_, *r*_12_, *r*_14_ (associated with *BioSD*^2^), in the order they appear. (d) The parameters increased or decreased in scenario (c) are now decreased or increased, respectively, in the same order. Simulation scenarios (a) and (b) correspond to perturbations within the network to be controlled while (c) and (d) to perturbations within the controllers’ architecture. For the latter two scenarios, PIDA-based simulations last longer due to the higher number of parameters being perturbed. Additionally, except for scenario (a), PID control eventually leads to closed-loop instability and loss of the RPA property, whereas PIDA control consistently performs as expected. Notably, in scenarios (c) and (d), the set-point changes resulting from perturbing *R*_*rs*_ and *r*_16_ (see Section IV) are always trackable by the PIDA controller, unlike with the PID controller.

**Fig. 5:**
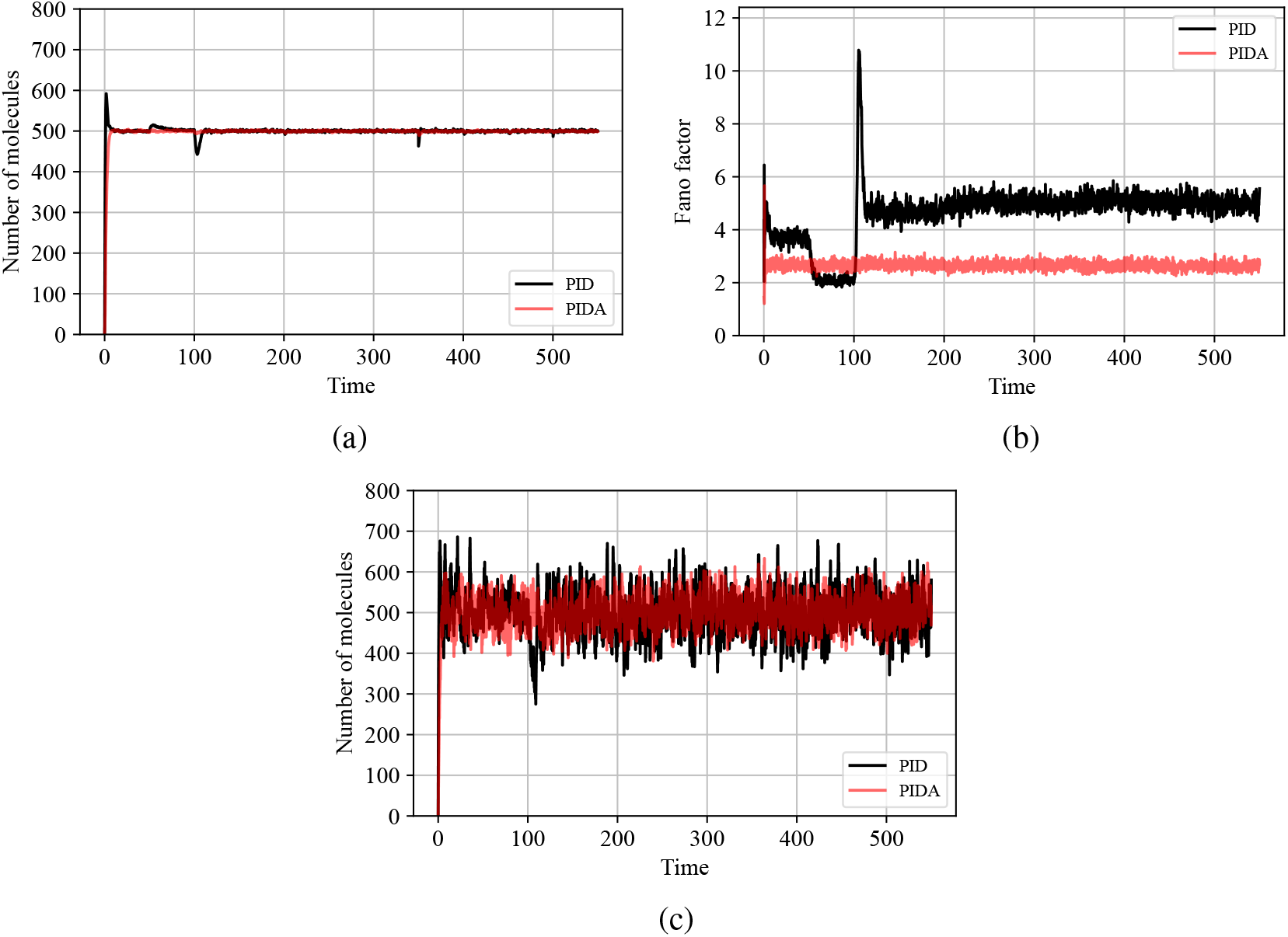
Considering the simulation scenario depicted in Fig. 4(a), the corresponding (b) stochastic means (c) Fano factors, *F*_*a*_(*t*), and (d) two (individual) stochastic trajectories (one for each case) are computed using SSA. The statistics presented in (a) and (b) are derived from 10^3^ stochastic trajectories. The conversion from concentration (in the deterministic setting) to the number of molecules (in the stochastic setting) was made as discussed in Fig. 3.

**Fig. 6:**
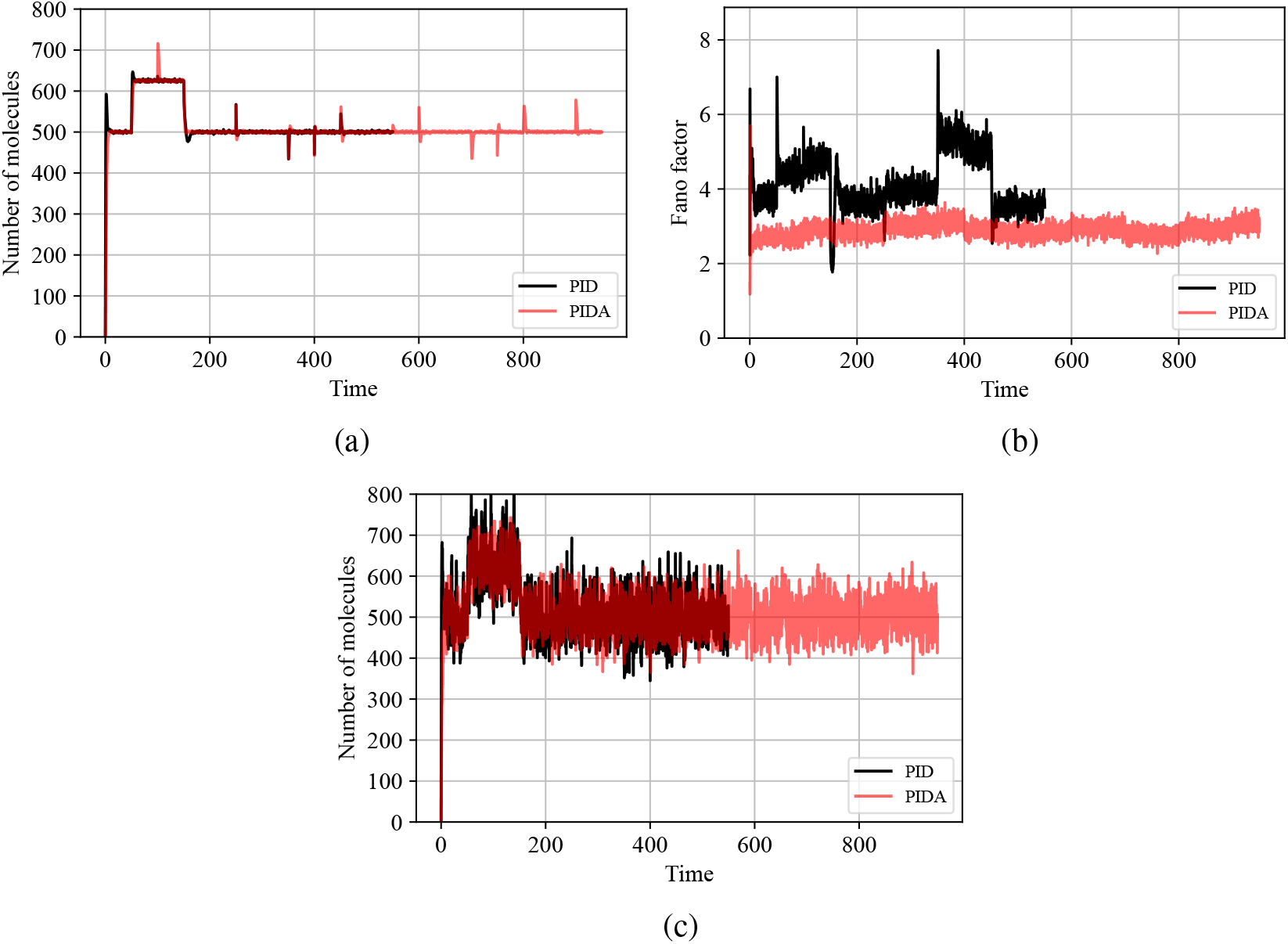
Considering the simulation scenario depicted in Fig. 4(c), the corresponding (b) stochastic means, (c) Fano factors, *F*_*a*_(*t*), and (d) two (individual) stochastic trajectories (one for each case) are computed using SSA. The statistics presented in (a) and (b) are derived from 10^3^ stochastic trajectories. The conversion from concentration (in the deterministic setting) to the number of molecules (in the stochastic setting) was made as discussed in Fig. 3. Note that the last parameter perturbation in the PID case, which leads to closed-loop instability in the deterministic setting (see Fig. 4), is not taken into account.

The above simulation example aims to underscore the beneficial role that acceleration action can play in molecular control design. Considering a case where PID control does not provide a desired closed-loop behavior, we show that the addition of acceleration feedback control alone (without any further structural or tuning modifications) is sufficient to greatly enhance it in both deterministic and stochastic domains. Nevertheless, this may not always be the case. It would not be surprising if, in molecular control settings with different characteristics - in terms of, for instance, the structure of the process to be controlled or the available parameter regimes - other (potentially simpler) architectures within this family of controllers could prove to be a more suitable choice.

## VI. Conclusion

This paper extends the concept of PID biocontrollers by presenting the first biochemical implementation of PIDA (which can be simplified to PDA) control scheme. Part of its architecture includes a novel biochemical device called *BioSD*^2^, which can function as a speed and acceleration biosensor attenuating, in parallel, disruptive high-frequencies with respect to an input molecular signal. Furthermore, we apply our PIDA strategy to *in-silico* regulation of a three-species biochemical process. In particular, we examine a scenario where PID control underperforms and demonstrate that the addition of acceleration action leads to a significantly smoother and more robust output response with reduced stochastic fluctuations while preserving RPA.

The molecular designs proposed herein might support future theory-guided experimental implementations. For instance, since we use mass action kinetics-based CRNs, they can be translated into *in vitro* biochemical systems via DNA strand displacement technology, as outlined in [10], [29]. It has also been experimentally established that, within this context, tuning of reaction rates over multiple orders of magnitude is possible [37]. For *in vivo* implementations, constructing such architectures within a single cell might be challenging due to their high structural complexity. Nevertheless, our molecular designs could potentially be useful for multicellular control purposes. With appropriate adaptation, the constituent parts of our control scheme could be distributed among different interacting cell populations within a microbial consortium. Notably, such approaches have already been investigated for simpler CRNs [35], [38], [39].

Another interesting aspect left for future work is to perform a “large signal analysis”, as the analytical results provided in this paper are obtained via linear perturbation analysis. It must be pointed out, however, that our theoretical analysis is validated through numerical simulations based on the corresponding nonlinear models, where the output of interest always starts at zero (far from its steady state). Finally, as previously emphasized, for simplicity, our controller is architecturally based on the principle that the actuation and sensing processes act directly on the output species. It would be interesting, though, to extend our design to more general cases where the actuation involves other species within the network being controlled.

